# Dual mechanisms of ictal high frequency oscillations in human rhythmic onset seizures

**DOI:** 10.1101/2020.04.07.030205

**Authors:** Elliot H. Smith, Edward M. Merricks, Jyun-You Liou, Camilla Casadei, Lucia Melloni, Thomas Thesen, Daniel Friedman, Werner Doyle, Ronald G. Emerson, Robert R. Goodman, Guy M. McKhann, Sameer A. Sheth, John D. Rolston, Catherine A. Schevon

## Abstract

High frequency oscillations (HFOs) recorded from intracranial electrodes during epileptiform discharges are a proposed biomarker of epileptic brain tissue and may also be useful for seizure forecasting, with mixed results. Despite such potential for HFOs, there is limited investigation into the spatial context of HFOs and their relationship to simultaneously recorded neuronal activity. We sought to further understand the biophysical underpinnings of ictal HFOs using unit recordings in the human neocortex and mesial temporal lobe during rhythmic onset seizures. We compare features of ictal discharges in both the seizure core and penumbra (spatial seizure domains defined by multiunit activity patterns). We report differences in spectral features, unit-local field potential coupling, and information theoretic characteristics of HFOs before and after local seizure invasion. Furthermore, we tie these timing-related differences to spatial domains of seizures, showing that penumbral discharges are widely distributed and less useful for seizure localization.

## INTRODUCTION

High frequency oscillations (HFOs) are discrete neural population events, typically >50 Hz, recorded from intracranial electrodes in patients with focal epilepsy (Bragin et al., 2002; Fried et al., 1999; Jirsch, 2006). Since they were first described in the 1990s, HFOs have been investigated for their ability to identify epileptic brain tissue and predict outcomes following surgical resection (reviewed in Höller et al., 2015; Jacobs et al., 2012). However, there has been limited success in translating these early results to achieve improved surgical outcomes.

More recently, attention has focused on identifying subclasses of HFOs with high predictive value (Fedele et al., 2017; Jacobs et al., 2018). These efforts have been based on spectral characteristics. Specifically, HFOs have been classified into ripples (∼80-200 Hz) and fast ripples (∼200-800 Hz) (Buzsaki et al., 1992). These two types of HFOs have been studied extensively in the rodent and human medial temporal lobe (Alvarado-Rojas et al., 2015; Foffani et al., 2007; Ibarz et al., 2010; Menendez de la Prida et al., 2006). In hippocampal slices with impaired inhibition, a single cell can induce population bursting with pathological fast ripples (Menendez de la Prida et al., 2006), whereas when inhibition is maintained, the same cells induce oscillations that restrain population activity in time (Bazelot et al., 2016; Schlingloff et al., 2014). These studies have shown that pathological bursting is characterized by reduced spike timing reliability across a population during a burst, which results in a broadening of the high frequency LFP spectrum and increased prevalence of fast ripples (Foffani et al., 2007). To date, there have been few studies aimed at identifying the cellular mechanisms of pathological HFO generation in the human neocortex, and the context of these mechanisms within the framework of seizure dynamics.

Previously, our group focused on ictal HFOs as a method of identifying cortex that has been recruited into a seizure, correlating these high-amplitude, phase-locked, broadband ripples with multiunit firing from nearby microelectrode arrays that exhibited signatures of recruitment into the seizure. That is, sites demonstrating a well-defined interictal to ictal transition (tonic firing of ictal wavefront, followed burst firing that is synchronized to the dominant rhythm of the seizure; Figure 1) as opposed to sites demonstrating heterogeneous, non-phase-locked firing. Since pathological phse-locked is more likely to produce a high gamma signature in macroelectrode recordings than is the case with heterogeneous non-phase-locked firing (Eissa et al., 2016; Ray and Maunsell, 2011), we focused on ictal HFOs beginning several seconds after seizure onset as a clinically accessible correlate of the specific neuronal firing pattern defining ictal recruitment (Weiss et al., 2013) and as a predictor of postoperative seizure control (Weiss et al., 2015). In these studies, we also showed that high gamma activity at seizure onset generally correlates poorly with evidence of seizure recruitment, and is a relatively poor predictor of seizure outcome when compared to current clinical standard assessments.

**Figure 1.**
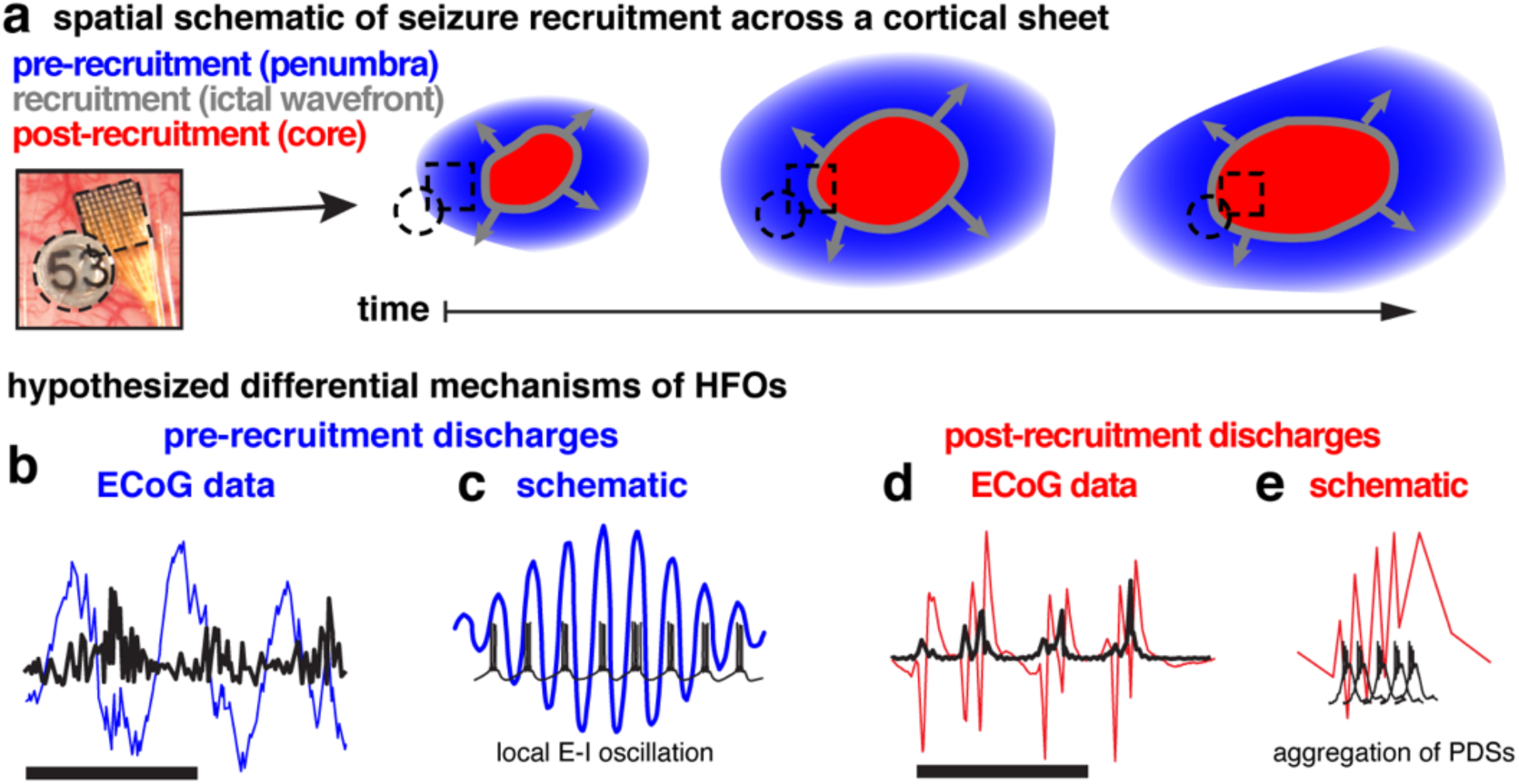
Schematics of spatial and temporal domains of focal seizures. A, Schematic of spatiotemporal patterns of seizure recruitment with color coded temporal labels and spatial labels in parentheses. Blue areas represent areas where pre-recruitment discharges occur (penumbra), the gray circles represent the slowly expanding ictal wavefront, and the red areas represent where post-recruitment discharges occur (ictal core). B, Example broadband pre-recruitment discharges (blue), and associated HFOs (black). Scale bar indicates one second of data. C, Schematic of hypothesized relationship between unit firing and pre-recruitment HFOs: unit firing is constrained in time by HFO phase during pre-recruitment discharges. D, Example broadband post-recruitment discharges (red), and associated HFOs (black). Scale bar indicates one second of data. E, Schematic of hypothesized relationship between unit firing and post-recruitment HFOs: unit firing consists of aggregations of paroxysmal depolarizing shifts.

However, this reasoning appears to be at odds with the long-held principle that electrophysiological changes at seizure onset are more likely to be indicators of epileptic tissue. To resolve this contradiction, we propose that pathological HFOs may arise from two contrasting mechanisms: paroxysmal depolarizing shifts resulting in spatiotemporally jittered population bursting that results dramatically increased *broadband high frequency* (BHF) LFP, and true narrowband oscillations arising from a combination of increased excitation and strong inhibitory interneuron firing, resulting in an exaggerated *narrowband gamma* rhythm. Such inhibitory-interneuron sculpted rhythms would be expected to occur in the ictal penumbra, i.e. at sites receiving strong synaptic currents generated from the seizure, yet exhibit the strong inhibitory restraint from surround inhibition preventing that area from being recruited into the ictal core. This narrowband gamma may occur at any time during a seizure, but may be most apparent at seizure initiation, when the ictal core is small relative to the area of surround inhibition that comprises the penumbra. To provide evidence for this hypothesis, we examine seizures characterized by both initial and delayed-onset HFOs at microelectrode-recorded sites, and contrast features of these two types of HFOs in the context of multiunit activity that is used both to define the timing and locations of seizure recruitment. We then translate these to clinically applicable measures, and leverage the resulting classification of HFOs to demonstrate differences in the ability of each type of HFO to localize seizures.

## RESULTS

### Temporal profile of seizure recruitment

Simultaneous microelectrode and standard clinical macroelectrode recordings were acquired in patients undergoing invasive EEG monitoring as part of neurosurgical treatment for pharmacoresistant focal epilepsy. There were 9 subjects with clinical grids and strip ECoG and “Utah” microelectrode arrays (MEA), and 16 subjects with stereo-EEG and Behnke-Fried depth microwire bundles. Recording was carried out continuously throughout the duration of the patients’ hospital stays in order to capture typical seizures. The seizures examined in this study were chosen based on the following criteria: 1) rhythmic slow oscillations at seizure onset, in a region that included at least one microelectrode recording site, 2) high quality action potentials (see Methods) recorded in the time periods leading into these seizures, and 3) multiunit activity (MUA) demonstrating the classic pattern of ictal recruitment, i.e. a period of tonic firing followed by repetitive burst firing (Schevon 2012, Merricks 2015). Seizures meeting these criteria allowed us to study epileptiform discharges locally before and after the passage of the ictal wavefront, that is when strong synaptic barrages affect an area of tissue versus when all neurons in a region exhibit strong rhythmic firing indicative of seizure recruitment. Six spontaneous seizures in four (three female) patients met these criteria, four seizures in the MEA group (two patients) and two seizures in the Behnke-Fried group (two patients), with stereotactically placed depth electrodes and microwires in mesial temporal lobe (MTL) structures. Clinical characteristics of these patients are provided in Table 1. A total of 322 multiunits during the 4 seizures were recorded from MEs (*mean* ± *std* = 80.5 ± 40.4 multi-units per seizure). The mesial temporal microwires yielded 7 multiunits between the two seizures. Example traces of broadband LFP, HFOs/BHF, and MUA recorded from a microelectrode are shown in Figures 2a,b, and c.

**Table 1.** Clinical characteristics of patients from whom seizures were examined.

| **Patient number**<br>**(age/sex)** | **1 (32/F)** | **2 (29/M)** | **3 (19/F)** | **4 (35/F)** |
| --- | --- | --- | --- | --- |
| **Type of Implant** | ECoG (UMA) | ECoG (UMA) | sEEG (B-F) | sEEG (B-F) |
| **Microwire array**<br>**implant location** | Left inferior<br>temporal gyrus,<br>2.5 cm from | Anterior middle<br>temporal gyrus | Mesial temporal<br>lobe (amygdala<br>& hippocampus) | Mesial temporal<br>lobe (amygdala<br>& hippocampus) |
| **Number of**<br>**seizures**<br>**examined** | 3 | 1 | 1 | 1 |
| **ILAE seizure**<br>**class** | Focal onset with<br>impaired<br>awareness | Focal to bilateral<br>tonic-clonic | Focal onset with<br>impaired<br>awareness | Focal onset with<br>impaired<br>awareness |
| **Clinically defined**<br>**seizure onset**<br>**zone** | Left basal<br>anterior<br>temporal | Right posterior<br>temporal with<br>secondary<br>generalization | Left mesial<br>temporal | Left mesial<br>temporal |
| **Resection Site** | Left basal<br>anterior<br>temporal | Right anterior<br>temporal with<br>inferior and<br>middle temporal<br>gyrus | Laser ablation of<br>left mesial<br>temporal lobe | No resection:<br>responsive<br>neurostimulator |
| **Surgical**<br>**outcome (follow-**<br>**up at time of**<br>**writing in**<br>**months)** | Engel 1a (55) | Engel 1a (36) | Engel 1d (32.7) | Engle 2c (17.4) |

**Figure 2.**
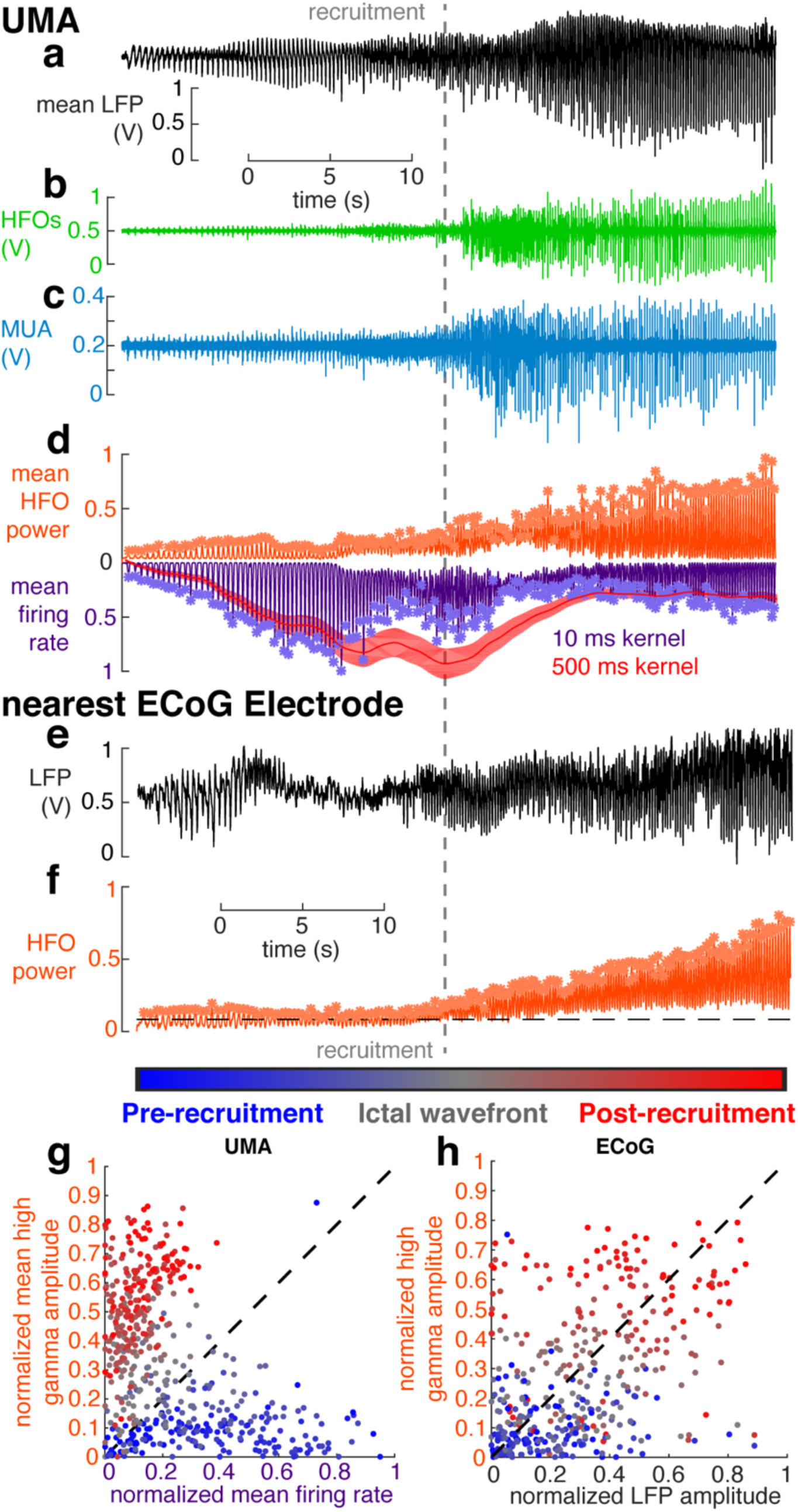
Temporal profile of seizure recruitment. The gray dotted line indicates the time of recruitment determined for MUA across the UMA. A, Mean LFP recorded across electrodes on the UMA. B, Example data from a single electrode on the UMA filtered in the HFO range, between 50 and 200 Hz. C, Example data from a single electrode on the UMA filtered in the MUA range, between 300 and 3000 Hz. D, Normalized mean MUA firing rate (purple), and HFO power (orange) for all channels on the UMA. Asterisks indicate detected discharges. The red trace shows the slow estimation of firing rate from which recruitment time was defined. E, broadband LPF from the nearest ECoG electrode to the UMA. F, Normalized HFO power on the same ECoG electrode as in e. Asterisks indicate detected discharges. G, Normalized MUA firing rate and HFO power for all detected discharges across 4 neocortical seizures plotted against each other and color-coded by when they occur in the seizure. H, Normalized LFP amplitude and HFO power for all detected discharges on the nearest ECoG electrode to the UMA across 4 neocortical seizures plotted against each other and color-coded by when they occur in the seizure.

The timing of ictal invasion of each microelectrode site was determined from the a transient pattern of continuous, asynchronous firing that is spatially organized into an ictal wavefront and is an indicator of inhibitory collapse (Figure 2d; Schevon et al., 2012a). This temporally divides the seizure, and thus its associated electrophysiology activity, into two stages: a *pre-recruitment* stage, in which the seizure has not yet invaded the microelectrode recording site, and a *post-recruitment* stage after seizure invasion (Smith et al., 2016). The pre-recruitment stage is the temporal analog of the penumbra, as described in our earlier reports (Schevon et al 2012). Consistent with our prior studies, the transition between the two stages could not be directly visualized in low frequency (< 50 Hz) recordings. Ictal discharges were detected by identifying BHF peaks (> 50 Hz; macroelectrodes; Figures 2e,f; Jirsch, 2006)) and MUA (microelectrodes; Figure 2d). Each detected discharge was matched between the two recording modalities and classified as either pre-recruitment or post-recruitment depending on timing relative to ictal wavefront passage recorded on the nearest microelectrode. In each seizure 287.5 ± 51.4 (mean ± s.d. per seizure) discharges were detected. The same number of pre and post-recruitment discharges were retained for further analysis (N = 154.8 ± 116.2 (mean ± s.d. per seizure)) and differences between neurophysiological features of these discharges were tested in a statistically balanced fashion.

Comparing the normalized mean amplitudes of MUA and BHF signals during each discharge yielded two discriminable axes for differentiating pre vs. post recruitment discharges. 74.4 % of pre-recruitment discharges had higher normalized MUA firing rates than normalized BHF amplitudes, and 94.5% of post recruitment discharges had higher normalized BHF amplitudes than normalized MUA firing rates. Accordingly, and as previously observed during seizures (Smith et al., 2016), normalized MUA firing rates and BHF amplitudes across all discharges were anticorrelated (Figure 2g; Pearson’s r = −0.44, N = 463, p < 10^−8^). Although MUA is not available from clinical ECoG or sEEG recordings, BHF amplitude differences still provided a relatively informative axis of discriminability between pre- and post-recruitment discharges. However, for macroelectrodes, the peak LFP amplitude and BHF amplitudes during discharges were significantly correlated (Figure 2h; Pearson’s r = 0.42, N = 400, p < 10^−6^).

### Differential MUA coupling to narrow-band HFOs before and after recruitment

We hypothesized that maintenance of inhibition is critical for generation of pre-recruitment HFOs, i.e. that an exaggerated expression of the feed-forward inhibition mechanism is associated with narrowband gamma oscillations, which have previously been associated with parvalbumin-positive interneurons (Freund and Katona, 2007; Isaacson and Scanziani, 2011; Roux and Buzsáki, 2015). We began by examining proposed multiunit indicators of maintained inhibition (Menendez de la Prida and Trevelyan, 2011). Specifically, we hypothesized that if inhibition were maintained during an epileptiform discharge, coupling of multiunit firing to the *dominant* LFP frequency (see methods for operational definition) of the surrounding population should exhibit phase symmetry and reliability, which would reflect a true narrobaend oscillation of excitatory and inhibitory drive. Alternatively, if inhibition has failed, multiunit events should couple to more asymmetrically-distributed phases of the dominant high frequency activity.

Our hypothesis was confirmed by the multiunit event coupling relationship between pre- and post-recruitment discharges, examples of which are shown in Figures 3a and 3b. Pre- and post-recruitment spike coupling distributions were significantly different (permutation Kuiper tests, all V_2000_ > 7.34, all p < 0.001). More specifically, neocortical unit firing during pre-recruitment discharges coupled to two specific and opposing phases of narrowband gamma oscillations, in a bimodal circular distribution (Figure 3c), however neocortical unit firing during post-recruitment discharges coupled more broadly to a single phase of gamma oscillations in a unimodal circular distribution (Figure 3d). The variance of this unimodal distribution was significantly greater than that of the angle-doubled bimodal distribution (two-sample tests for equal concentration parameters, all p < 0.007, all F > 1.03). Critically, these differences were only determined by examining the dominant, patient-specific high-frequency bands. If pre-defined narrowband gamma (30-60 Hz) and ripple (70-200 Hz) frequency ranges were used, no significant differences in unit coupling between pre- and post-recruitment discharges were observed (permutation Kuiper tests, all V_2000_ < 1.91, p > 0.36 for low gamma; permutation Kuiper tests, all V_2000_ < 1.25, p > 0.46, for high gamma).

**Figure 3.**
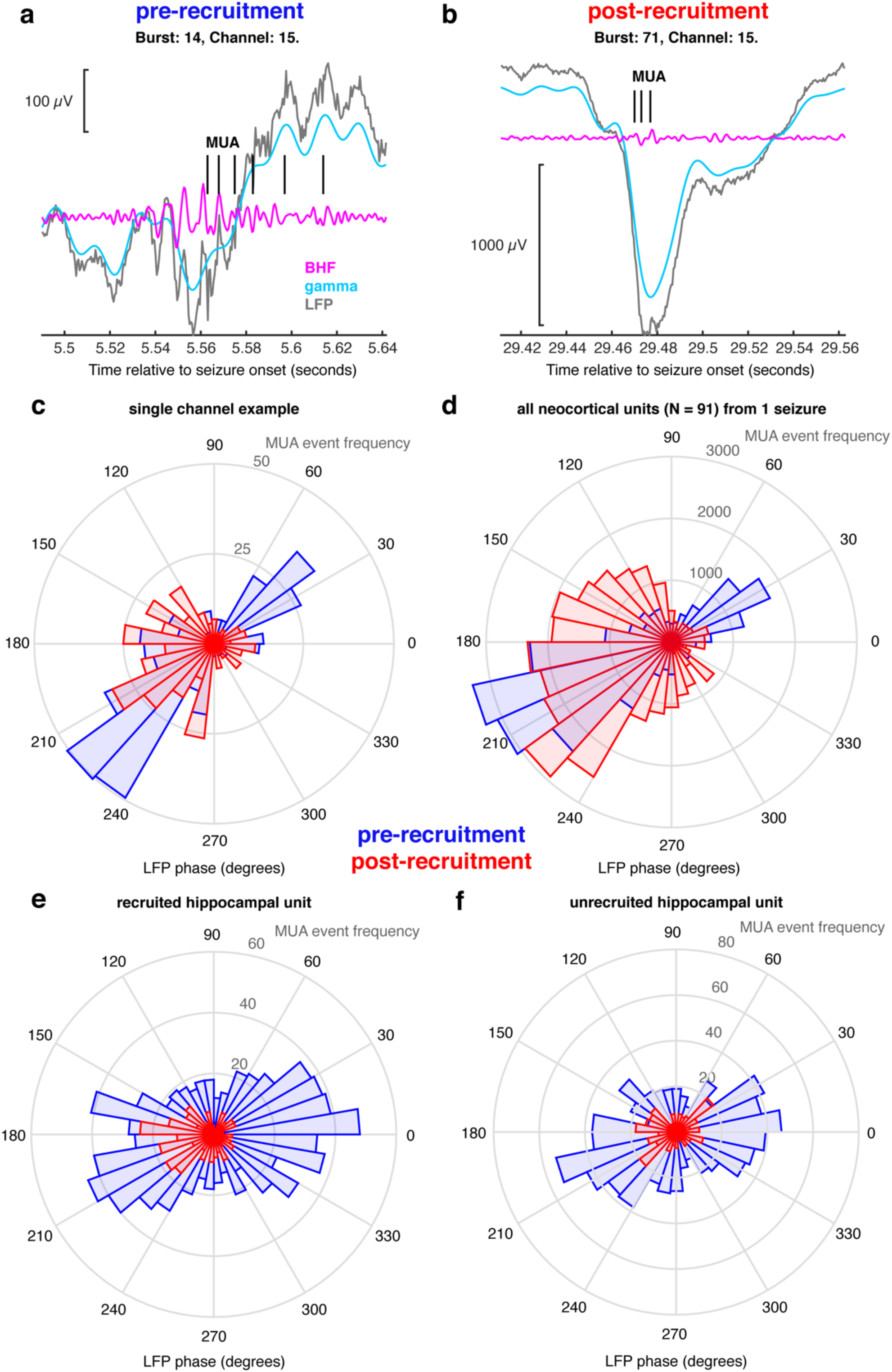
Multiunit coupling to HFOs. A, a single example pre-recruitment discharge recorded on the UMA showing broadband LFP (gray), filtered in the gamma range (cyan) and BHF range (magenta). MUA event times are shown in black. B, an example post-recruitment discharge, showing the same signals color coded in the same way as in A. C, A circular histogram counting multiunit event times for every 10 degrees of phase of the dominant frequency. Blue bins show coupling during pre-recruitment discharges, and red bins show coupling during post-recruitment discharges. D, A circular histogram counting all multiunit event times in a single seizure for every 10 degrees of phase of the dominant frequency. Blue bins show coupling during pre-recruitment discharges, and red bins show coupling during post-recruitment discharges. E, An example hippocampal unit that was recruited into the seizure. Blue bins show coupling during pre-recruitment discharges, and red bins show coupling during post-recruitment discharges. F, An example hippocampal unit that was not recruited into the seizure. Blue bins show coupling during pre-recruitment discharges, and red bins show coupling during post-recruitment discharges.

MTL units exhibited similar dominant frequency phase-locking as that for neocortical units. Figure 3f shows multiunit event coupling from an MTL unit that was recruited into the seizure, demonstrating a shift in phase-locking between pre- and post-recruitment events (permutation Kuiper test, V_200_ = 6.8, p < 0.001). However MTL units that were not recruited into the seizure (n = 6) maintained the bimodal spike-timing distribution during post-recruitment discharges (Figure 3f; permutation Kuiper test, V_200_ < 2.16, p > 0.61). Not only do these data provide evidence for differential unit coupling in different spatial domains in the same seizure, these results show that both neocortical and MTL multiunit events couple to the dominant narrowband gamma frequency of pre-recruitment discharges, with a shift in multiunit event coupling occurring in post-recruitment discharges. This provides support for the hypothesis that pre-recruitment discharges occur in tissue in which feedforward inhibition is maintained, and that this mechanism is similar in both hippocampus and neocortex.

### Spatiotemporal profile of seizure recruitment across clinical recordings

As schematized in Figure 1a, and described in the previous sections, neuronal data from microelectrodes indicate a spatiotemporal structure of seizure expansion, where adjacent groups of cells are successively recruited into the ictal core. We use the terms pre- and post-recruitment to define temporal seizure domains before and after passage of the ictal wavefront. We use the terms ictal penumbra, and ictal core, to describe the corresponding spatial domains (Figure 1a). Even as this successive recruitment is occurring, the effects of the seizure core extend beyond the ictal wavefront, to the penumbra, where inhibition is likely preserved, yet neural ensembles are assailed by discharges from the adjacent, already recruited tissue. In the previous section we showed data indicating that the high frequency LFP during discharges in these two territories has different mechanisms. In the following two sections, we examined correlates of these two types of discharges on a larger, more clinically relevant scale, relating changes in the high frequency LFP on clinical macroelectrodes to intense, disorganized, phase-locked action potential firing in the seizure core.

In order to extend our analysis to HFOs recorded on macroelectrodes throughout the brain, we utilized phase-locked BHF, a previously defined ECoG-based measure of recruitment (Weiss et al., 2015, 2013), to operationally define the spatial extent of ictal recruitment across the entire coverage area. We maintained the same temporal delineation of recruitment defined by passage of the ictal wavefront on simultaneously recorded microelectrodes. Examples of recruited and unrecruited macroelectrodes are shown in Figures 4a,b,c, and d. Phase-locked BHF yielded clearly bimodal distributions of channels during the post-recruitment phase (Figure 4e; t_363_ = 2.8, p = 0.006), indicating that not all implanted macroelectrodes were recruited.

**Figure 4.**
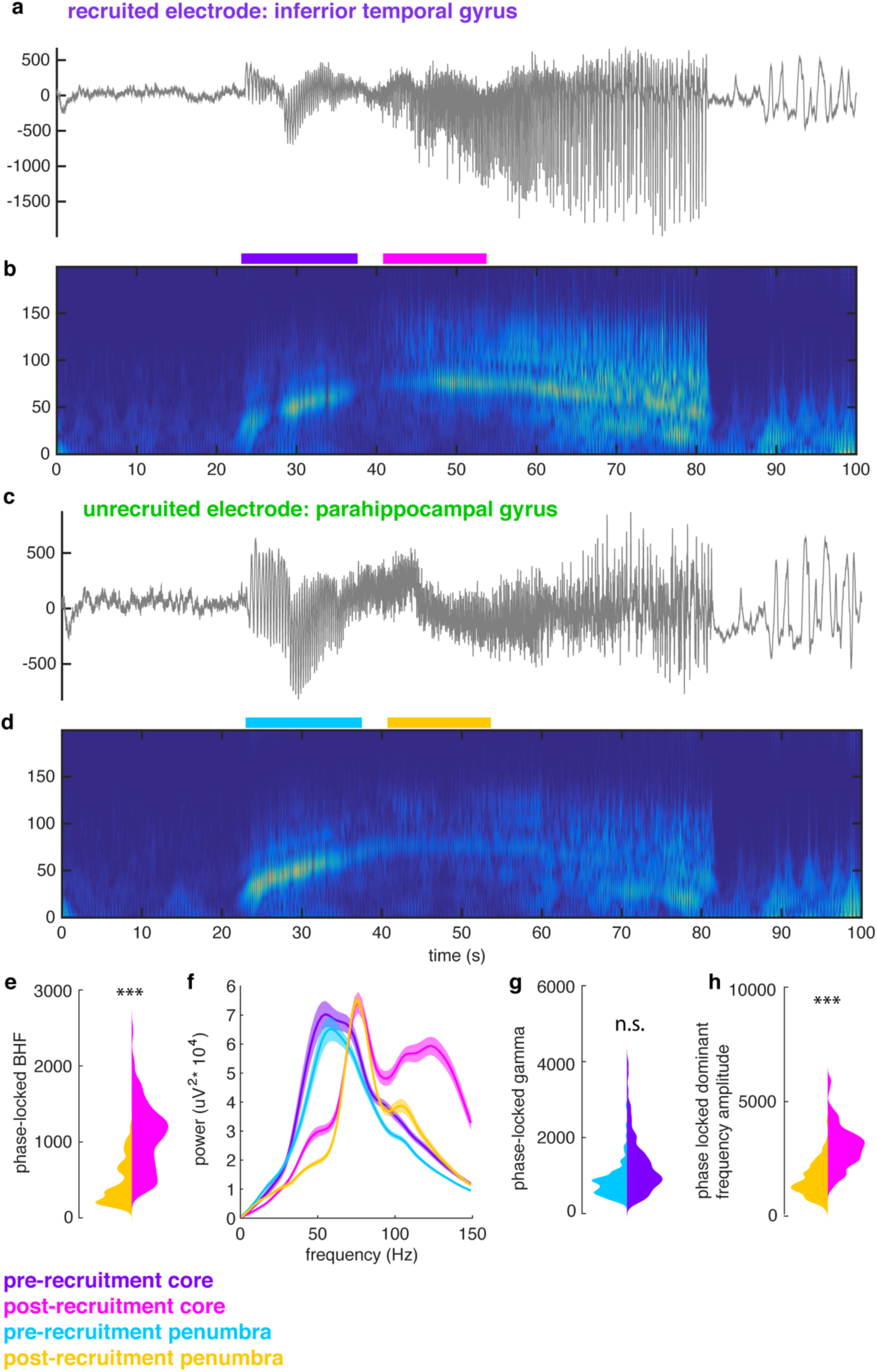
Spatiotemporal profile of recruitment across clinical electrodes. A, example broadband LFP from an ECoG electrode that is recruited into the ictal core. B, spectrogram of the seizure recorded on the electrode shown in A. C, example broadband LFP from an ECoG electrode that remains in the penumbra. D, spectrogram of the seizure recorded on the electrode shown in C. E, violin plots showing the distributions for post-recruitment phase locked BHF (95 – 200 Hz) in the core (magenta) and the penumbra (orange). Asterisks indicate significant difference between the two distributions in the mixed effects model F, Mean spectra for all electrodes in the core and penumbra electrode groups for the pre- and post-recruitment periods. These epochs are shown as colored bars in B and D. G, violin plots showing the distributions for pre-recruitment phase-locked narrowband gamma (30 – 70 Hz) in the core (purple) and the penumbra (cyan). n.s. indicates no significant difference in the mixed effects model. H, violin plots showing the distributions for post-recruitment phase locked dominant frequency (approximately 90 Hz) in the core (magenta) and the penumbra (orange). Asterisks indicate significant difference between the two distributions in the mixed effects model.

We therefore classified discharges recorded on macroelectrodes into 4 mutually exclusive categories that correspond to different *spatiotemporal* seizure domains: pre-recruitment discharges on core macroelectrodes, post-recruitment discharges on core macroelectrodes, pre-recruitment discharges on penumbral macroelectrodes, and post-recruitment discharges on penumbral macroelectrodes (i.e. discharges after seizure recruitment on the microelectrode array in contacts that are never recruited into the ictal core). That is, in the *temporal* domain, pre- and post-recruitment discharges were distinguished based on the MUA-based temporal designation described above. In the *spatial* domain, core and penumbra contacts were distinguished by clustering the bimodally-distributed post-recruitment phase-locked high gamma values into the two distinct unimodal components of the overall distributions using Gaussian Mixture Models. The mean spectra for these four distributions are shown in Figure 4f. The pink spectrum clearly shows how the post-recruitment core is distinguished from the other spatiotemporal categories by its increased BHF.

As the tissue under the penumbra electrodes is not recruited, we compared penumbral discharges to discharges located within the core, occurring both before and after the ictal wavefront, in order to understand large-scale properties of the penumbra. We found that during the pre-recruitment period, phase-locked narrowband gamma did not differ between core and penumbral macroelectrodes (Figure 4g; t_363_ = 0.2, p = 0.82). Examining the Bayes Factor from this model suggested that narrowband gamma oscillations occurring in the pre-recruitment period do not distinguish the core and penumbra (Bayes Factor = 1.2*10^−32^). Unsurprisingly, this similarity was also evident between the phase-locked dominant frequency LFP (largely narrowband gamma) in the core and penumbra during the pre-recruitment period (t_363_ = 1.9, p = 0.05). However, phase-locked dominant frequency power during post-recruitment discharges (largely BHF) differed between core and penumbral electrodes (Figure 4h; t_363_ = 8.5, p < 0.01). Accordingly, dominant frequencies of pre-recruitment discharges were lower than those from the post-recruitment epoch (pre-recruitment mean ± std: 57.94 ± 11.38; post-recruitment mean ± std: 65.01 ± 16.95; t_728_ = −3.3, p = 0.001). Together these results show that the pre-recruitment time period and penumbral spatial domain are marked by a narrowband gamma oscillation whose dominant frequency increases until the point of seizure recruitment. Importantly, this narrowband gamma oscillation does not differentiate between core and penumbra territories, as it can appear across widely distributed brain areas including those distant from the seizure onset zone. Given that all but one of the patients were seizure free after resection surgery (apart from the patient who received responsive neurostimulation, with more than 33 months follow-up at time of writing), this provides additional evidence that pre-recruitment HFOs are not reliable biomarkers of epileptic brain in rhythmic onset seizures. Following recruitment, BHF increases specifically in the slowly expanding seizure core.

### Gauging seizure recruitment with discharge complexity

We next sought to further clarify the features underlying differences between pre- and post-recruitment discharges, and relate these differences to the pathological nature of HFOs previously reported in animal models (Foffani et al., 2007; Ibarz et al., 2010; Valero et al., 2015). The most obvious qualitative difference between HFOs during pre- and post-recruitment discharges was that HFOs during pre-recruitment discharges appeared smoother and less complicated than those during post-recruitment discharges. We sought to quantify this feature by measuring entropy of the relative distributions of recorded voltages through the duration of each discharge (differential entropy) and of the relative distribution of LFP frequencies (spectral entropy) during each discharge. These two quantities may alternatively be thought of as respective measures of time-domain and spectral-domain complexity of each epileptiform discharge. As in the aforementioned studies, we compare these measures to the proportion of the spectrum in the ripple and fast ripple ranges (ripple and fast ripple indices; see methods).

Figures 5a,b show the distributions of the fast ripple indices and spectral entropies for all discharges and MEA electrodes for a single seizure. After accounting for any random variance that may have occurred across patients and seizures using linear mixed-effects models, we found that pre- and post-recruitment discharges recorded on microelectrodes had no significant difference in fast ripple indices (Figure 5c; generalized linear mixed effects model: index ∼ 1 + epoch + (1 + epoch | seizure); t_87,723_ = −0.59, p = 0.55; Bayes factor = 0) or differential entropy (Figure 5d; generalized linear mixed effects model: entropy ∼ 1 + epoch + (1 + epoch | seizure), t_87,723_ = 0.005, p = 0.99; Bayes factor = 3.5*10^−120^). However, spectral entropies were significantly higher for post-recruitment discharges (Figure 5e; generalized linear mixed effects model: entropy ∼ 1 + epoch + (1 + epoch | seizure), t_87,723_ = −3.09, p < 0.002).

**Figure 5.**
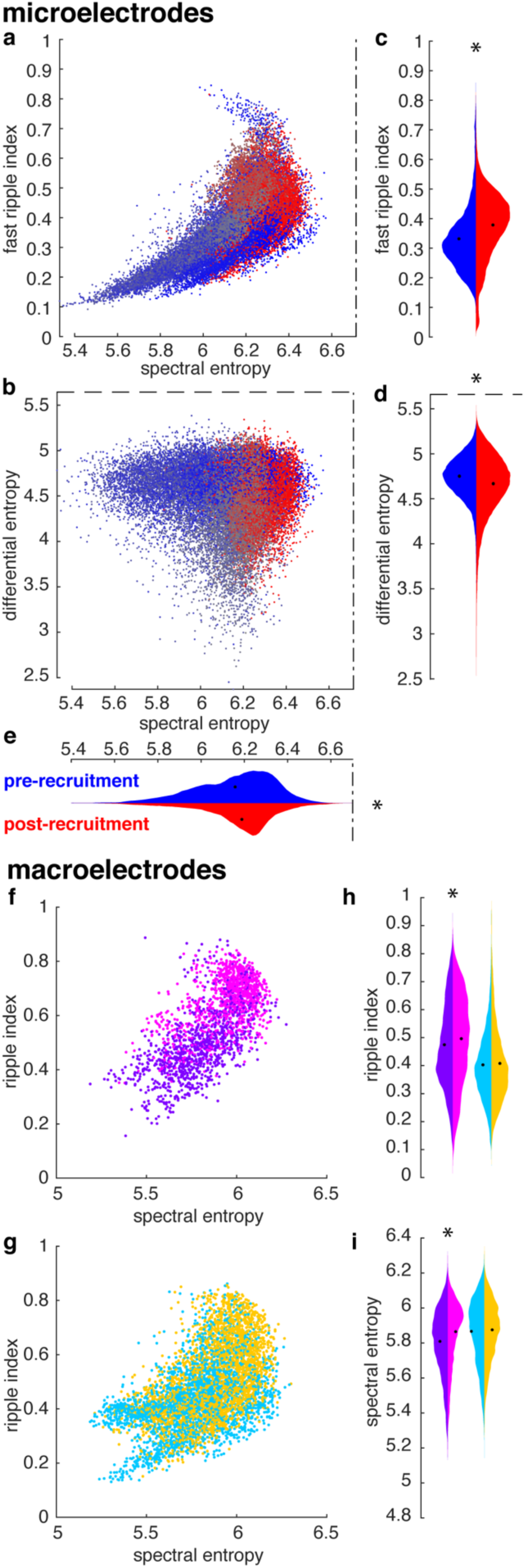
gauging seizure recruitment by discharge complexity. A, spectral entropy plotted against the fast ripple index for all pre and post –recruitment discharges recorded on the UMA. Points are colored as in the color bar in figure 2. Dash-dot lines indicate the theoretical limit for spectral entropy. B, Spectral entropy plotted against differential entropy for all pre and post –recruitment discharges recorded on the UMA. Points are colored as in the color bar in figure 2. Dash-dot lines indicate the theoretical limit for spectral entropy. Dashed lines indicate the theoretical limit for differential entropy. C, Violin plots of distributions of fast ripple indices across all discharges seizures recorded on UMAs for pre and post –recruitment discharges shown in blue and red, respectively. D, Violin plots of distributions of differential entropy across all discharges seizures recorded on UMAs for pre and post –recruitment discharges shown in blue and red, respectively. Dashed lines indicate the theoretical limit. E, Violin plots of distributions of spectral entropy across all discharges seizures recorded on UMAs for pre and post –recruitment discharges shown in blue and red, respectively. Dash-dot lines indicate the theoretical limit. F, spectral entropy plotted against the ripple index for all ECoG electrodes that get recruited into the core. Dots are color-coded as in figure 4. G, spectral entropy plotted against the ripple index for all ECoG electrodes that do not get recruited into the core. Dots are color-coded as in figure 4. H, Violin plots of distributions of ripple indices recorded on ECoG across all seizures for pre and post –recruitment discharges for the core and penumbra. Distributions are color-coded as in figure 4. I, Violin plots of distributions of spectral entropy recorded on ECoG across all seizures for pre and post –recruitment discharges for the core and penumbra. Distributions are color-coded as in figure 4.

In order to determine whether these measures translate to clinical ECoG recordings, we also calculated similar entropy measures for discharges recorded on macroelectrodes, again employing the four spatiotemporal seizure domains described above. The sampling rates typically used to record clinical data, and that were used to record these data, do not allow for the detection of fast ripples, but a wealth of evidence, both previously presented, and presented herein, suggest that broadband, high frequency LFP is informative about the intense firing of recruited units (Weiss et al., 2013). We therefore calculated a “ripple index” (see Methods for operational definition) analogous to fast ripple indices derived from microelecrode recordings and to compare with spectral and differential entropy measures derived from the signals recorded on macroelectrodes. This index differs from standard HFO definitions by its basis on the area under the normalized spectrum, rather than amplitude of particular spectral bands. The “ripple index” therefore incorporates widening of the spectrum, as well as the normalized amplitude of the ripple band, relative to the rest of the spectrum. Figures 5f and 5g show the distributions of the “ripple index” and spectral entropy for each discharge in a representative seizure. This measure was significantly higher for post-recruitment discharges in the putative ictal core (Figure 5h; generalized linear mixed effects model: index ∼ 1 + epoch + (1 + epoch | seizure), t_28,418_ = −2.23, p = 0.02; Weiss et al., 2013), yet ripple indices were not significantly different for pre- and post-recruitment discharges in the penumbra (Figure 5h; generalized linear mixed effects model: index ∼ 1 + epoch + (1 + epoch | seizure), t_26,706_ = −1.4, p = 0.14). Similar to the microelectrode results, there was neither a significant difference in differential entropy between pre- and post-recruitment discharges in the core (generalized linear mixed effects model: entropy ∼ 1 + epoch + (1 + epoch | seizure), t_28,418_ = 0.52, p = 0.6) nor the penumbra (generalized linear mixed effects model: index ∼ 1 + epoch + (1 + epoch | seizure), t_26,706_ = 0.08, p = 0.94). However, there was a significant difference in spectral entropy between pre- and post-recruitment discharges in the ictal core (Figure 5i; generalized linear mixed effects model: index ∼ 1 + epoch + (1 + epoch | seizure), t_28,418_ = 3.8, p = 0.0001), yet not in the penumbra (Figure 5i; generalized linear mixed effects model: index ∼ 1 + epoch + (1 + epoch | seizure), t_26,706_ = 1.7, p = 0.07). These results further confirm, that increased complexity and significant broadening of the LFP spectrum in the ictal core is related to large phase-locked increases in BHF, and disorganized, intense unit firing that localizes specifically to the ictal core.

## DISCUSSION

Here we provide evidence that early discharges in spontaneous human seizures with rhythmic onset morphology occur in brain tissue in which feed-forward inhibition is maintained. More specifically, we show that discharges occurring before the neuronal signature of recruitment on average exhibit phase-locked narrowband gamma-range activity (approximately 30 – 80 Hz) across a broad cortical area. Multiunit firing couples to these narrowband gamma oscillations in a bimodal pattern during pre-recruitment discharges. Following recruitment, a more restricted cortical area that corresponds to the seizure core exhibits a high-amplitude BHF activity that is phase locked to the seizure’s dominant rhythm. This BHF activity is tightly temporally linked to phase-locked multiunit activity, reflective of actively seizing brain tissue. These post-recruitment discharges exhibit increased complexity on both microelectrodes and clinical electrodes. We therefore translate our understanding of the dynamics of multiunit activity during seizures to clinical recordings with important implications for seizure localization in the clinical setting, and demonstrate that early discharges in rhythmic onset seizures exhibit marked narrowband gamma, and that these are present outside of seizing brain tissue.

Together these results provide further means to decode the neuronal and synaptic processes underlying the discharges that make up focal seizures, and therefore have important clinical implications. While pre-recruitment discharges indicate seizure activity somewhere, they are produced by an inhibitory response to topologically adjacent seizing tissue. That is, inhibitory firing sculpts excitatory bursts into a fast oscillatory structure, resulting in a narrowband gamma oscillation. In the seizure core, dominated by paroxysmal depolarizing shifts, this sculpting effect is absent, and the appearance of a pseudo-oscillation, termed BHF, arises from summated, jittered postsynaptic activity (Eissa et al., 2016). Importantly, early phase-locked narrowband gamma oscillations do not necessarily precede post-recruitment phase-locked BHF. These features of epileptiform discharges could however be extremely informative if interpreted as follows: early discharges with increased phase-locked narrowband oscillatory gamma power are likely not yet recruited into the seizure core, yet they are receiving strong excitatory synaptic input from a connected cortical territory. We speculate that an analogous situation may occur with interictal epileptiform discharges, enabling more accurate subclassification of pathological interictal HFOs, given that surround inhibition is known to operate during these events as well.

The current results also inform our basic understanding of seizure pathophysiology. By replicating results from the animal literature in which destructive, pharmacological, or optogenetic manipulations have been performed. Previous studies in animal models have shown that neuronal firing is less precise when feed-forward inhibition is altered (Foffani et al., 2007; Ibarz et al., 2010; Miri et al., 2018; Trevelyan, 2009). This lack of action potential timing reliability has been related to several cellular processes. Changes in coupling to the high frequency LFP, as we report here, has been associated with cellular chloride loading in early ictal discharges (Alfonsa et al., 2015). However, in Alfonsa et al. such chloride loading was insufficient to induce seizure-like activity alone (Alfonsa et al., 2015). Both AMPA and acetylcholine agonists were able to rescue spike timing reliability, and blocking potassium channels increased spike timing reliability in hippocampal slices (Foffani et al., 2007). The current results support a combination of these schemas, suggesting that strong feedforward inhibition, likely mediated by parvalbumin-expressing GABAergic interneurons, is reflected in gamma range neuronal synchrony, which is unrelated, both spatially and temporally as we show here, to the BHF increases that correlate with unconstrained pyramidal cell firing and PDS (Jefferys et al., 2012).

As with physiological high frequency LFP, the generating mechanisms of HFOs in human epilepsy remain an open area of investigation. Results from animal studies suggest jittered spike timing as a mechanism for converting ripples to fast ripples, or broadening the high frequency spectrum in general, both in hippocampus (Bragin et al., 2002; Foffani et al., 2007; Ibarz et al., 2010) and neocortex (Alfonsa et al., 2015). Accordingly, we propose that HFOs associated with PDS, while meeting commonly accepted HFO criteria, are not true oscillations, but rather are generated from summated spatiotemporally jittered fast potentials, which are far stronger during PDS than during non-pathological neuronal bursting activity (Eissa et al., 2016). Such strong, temporally disorganized neuronal firing results in an aperiodic shift in the high frequency spectrum, resulting in broadband high frequency activity (BHF) (Manning et al., 2009; Miller, 2010). BHF contrasts with HFOs that may be regarded as true oscillations, such as those produced from phasic population firing that exhibit clear, periodic excursions from the overall (≈ *f*^−*n*^) shape of the neural spectrum (Buzsáki et al., 2012; Haller et al., 2018). It is hypothesized that synchronized fast-spiking inhibitory interneuronal activity plays the crucial role of delineating temporal windows for pyramidal cells to fire (Menendez de la Prida and Trevelyan, 2011). Ironically, such true oscillatory HFOs may therefore occur in brain sites that have not been invaded by seizures due to an intact inhibitory restraint mechanism (Schevon et al., 2012b; Trevelyan et al., 2007). We therefore propose that there are two distinct mechanisms for pathological HFOs, that identify two HFO classes with differing utilities as biomarkers.

## MATERIALS AND METHODS

### Subjects and ethics statement

Four patients with pharmacoresistant epilepsy were included in this study. Despite these relatively small sample sizes of patients and seizures, our statistics are based on the numbers of units, electrodes, and discharges observed in these seizures and are therefore sufficiently powered. The included patients were implanted with standard clinical ECoG electrodes (2 patients) or stereo electroencephalographic (sEEG) electrodes as part of the monitoring prescribed for surgical treatment of their epilepsy. In addition, all four patients were implanted with microelectrodes. ECoG patients received a “Utah-style” microelectrode array (MEA), and stereo EEG patients had Behnke-Fried (B-F) style microelectrodes. MEAs are 4 × 4 mm square arrays with 96 recording microelectrodes (10 × 10 electrodes, with the corner electrodes disconnected) that penetrate 1 mm into layer 4/5 of human association cortex (Schevon et al., 2012b). B-F microelectrodes consist of a bundle of eight microelectrodes that protruded approximately 4 mm from the tips of depth electrodes. Detailed methodology for surgical implantation of these arrays are described in (House et al., 2006) for MEAs and (Misra et al., 2014) for B-Fs. The MEAs were all implanted into the clinically defined seizure onset zone. Both the New York University and Columbia University Medical Center Institutional Review Boards approved this study and each patient provided informed consent prior to surgery.

### Data collection and preprocessing

Neural data were acquired from each microelectrode at 30 kilosamples per second (0.3 Hz– 7.5 kHz bandpass, 16-bit precision, range ± 8 mV) using a Neuroport Neural Signal Processor (Blackrock Microsystems, LLC). ECoG and sEEG signals were acquired using the clinical amplifier (Natus Medical, Inc.) at either 500 or 2000 samples per second (0.5 Hz high pass, low pass set to 1/4 sampling rate, 24-bit precision). All ECoG data with sampling rates higher than 500 samples per second were subsequently downsampled to 500 samples per second in order to facilitate comparisons across patients and clinically-relevant sampling rates.

### Spectral analysis

Spectrograms of full seizures and of individual discharges were each generated using 5-cycle Morlet wavelet scaleograms using 107 logarithmically-spaced frequency bins from 1 to 200 Hz for ECoG data, and 1 to 850 Hz for MEA data. The absolute values of these scaleograms were normalized by a theoretical spectrum, *S*, such that

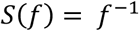

Where *f* represents the frequency range of the spectrum. These methods were used for both the full duration of the seizure and for each discharge and for both ECoG and MEA recordings.

### Dominant frequency analysis

In order to detect the dominant frequencies of seizure-associated LFP through the duration of the seizure, the aforementioned spectral analysis was carried out on each channel of electrophysiological data from 5 seconds before any discharge detection to 5 seconds after the final discharge detection. The maximum frequency in each time bin was then smoothed by a moving average with a width of one eighth of the voltage signal’s sampling rate. The resultant time series operationally defined the dominant frequency through the duration of each seizure.

### Per-discharge analyses

In order to understand differences among ictal discharges before recruitment into the ictal core, we first detected each ictal discharge using a thresholding and clustering process. First, all local maxima above one standard deviation in the mean LFP power between 50 and 200 Hz were detected. This method yielded a few false positives due to the noisy nature of LFP, particularly on ECoG. In order to remove false positives, the differences in amplitude and timing of the aforementioned detections were clustered into seven groups using k-means and outlying clusters were removed in a supervised manor until aberrant detections were no longer present. The remaining detection times were used for all further ECoG analyses.

We detected discharges on the MEA using a different procedure, which capitalized on the lack of noise in the MEA recordings and match firing rate and LFP measurements across discharges. Specifically, we detected discharges on the Utah array by cross-referencing the normalized maxima of firing rate and instantaneous high gamma amplitude over channels. First, both instantaneous high gamma and firing rate were estimated for each channel and convolved with a Gaussian kernel with 10 ms standard deviation. These signals were normalized by subtraction of their absolute minimum and division by their absolute maximum. Local maxima were detected in both high gamma and firing rate using differentiation. Each maximum exceeding one standard deviation of the firing rate or high gamma amplitude during the seizure was retained as an ictal discharge. Any discharge that occurred within 50 ms of the discharge peak was excluded from further analysis. Discharges that were detected in the BHF signal, but not in the firing rate signal, or vice versa, were also discarded. The resulting data are therefore comprised of co-occuring ictal discharges in the HFA and firing rate signals.

### MUA detection and time of recruitment

Since we previously showed that sorting single units during the recruited period is impossible, here we examine multiunit activity recorded on MEAs and microwires. MUA on these electrodes was determined by first zero-phase-lag filtering the data between 300 and 3000 Hz and then finding peaks greater than 3.5 times two thirds of the median absolute value of the filtered data (Merricks et al., 2015). The times of these peaks were retained as multiunit event times.

We determined that a multi-unit was recruited into a seizure from its pattern of firing. If the multi-unit exhibited a period of intense tonic firing that was flanked in time by discharging, we considered that multi-unit as recruited. All of the multi-units from the MEA recordings were recruited into the seizure core. Only a single MTL unit from the microwires was recruited into the core. The time of recruitment was detected using methods from (Smith et al., 2016). Specifically, a Gaussian kernel with 500 ms width was convolved with the MUA on each channel in order to emphasize slow elements of the signal. The mean peak of this slow firing rate estimate was defined as the time of seizure recruitment.

### Action potential phase coupling

Action potential coupling was determined by extracting the phase from the complex elements of the analytical signal of the bandpass filtered LFP (96^th^ order fir filter) at ± 5 Hz above and below the dominant frequency of the LFP, calculated as aforementioned, for each MUA event time. Histograms of action potential times relative to LFP phase (36 bins) were then constructed for each unit and across units for each seizure. We tested for differences in coupling for each seizure using a modified Kuiper test in which a random partition of 2000 action potential times was taken from the cumulative density functions calculated above for both the pre and post –recruitment periods. The test statistic, *V*, was defined as

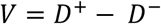

where *D*^*+*^ and *D*^*-*^ represent the maximum two-tailed distances between the cumulative density functions, as in a standard Kuiper or Kolmogorov-Smirnov test. This procedure was carried out 10,000 times and the test statistic calculated on data from all units in each seizure was compared to the permutation distribution in order to derive a p-value. Kuiper tests in which the test statistic controlled for sample size were carried out on a per-unit basis for MTL neurons using a permutation size of 200 (Koziol J. A., 1996).

### Defining recruited clinical electrodes using phase-locked measures

As a formal method for defining recruited vs unrecruited electrodes, Gaussian Mixture Models with two univariate normal density components were considered. These models were fit using the iterative expectation maximization algorithm. Each of the models converged and exhibited clearly-separated posterior probability distributions.

### Per-discharge entropy

In order to quantify the temporal complexity of voltages measured during ictal oscillations, *V(t)*, differential entropy of the signals, *h(V)*, for each discharge was calculated as

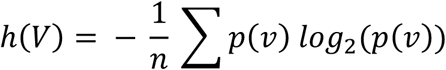

Where *p(v)* is the probability density function of voltage values for each discharge and *n* is the number of samples examined for each discharge, where *n* was 98 samples for macroelectrodes and 296 samples for microelectrodes.

The entropy of the signals in the spectral domain, as in (Foffani et al., 2007; Ibarz et al., 2010; Valero et al., 2015), were also examined for each ictal discharge recorded on the MEA. The spectral entropy was calculated as

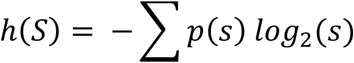

Where *p(s)* is the distribution of power in each frequency between 100 and 800 Hz. This measure has a maximum at the log of the number of frequency bins used in the spectral estimation. The current study used 50 frequency bins for ECoG and sEEG electrodes and 107 frequency bins for microelectrodes, yielding maximal entropies of 5.64 and 6.71 bits, respectively. This entropy measure was compared with spectral mode and fast ripple index, as calculated in (Ibarz et al., 2010). The spectral mode is the most prevalent frequency in the spectrum between 100 and 800 Hz and the fast ripple index is the proportion of the normalized spectrum above 250 Hz. We added an additional index to these analyses in order to observe similar relationships in macroelectrode recordings. This “ripple index” was defined as the proportion of the normalized spectrum between 80 and 200 Hz in spectra spanning frequencies from 1 to 250 Hz.

### Bayesian Statistics

Several of the linear mixed effects models (LMM) implemented did not exhibit significant slopes. We therefore sought to quantify the extent to which our beliefs about the hypotheses tested in these models were updated by using Bayesian analysis. We calculated Bayes factors for the LMM fixed effects from the models’ Bayesian Information Criteria (BIC) (Jarosz and Wiley, 2014). The Bayes Factors were calculated as

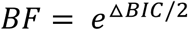

Where Δ*BIC* refers to the difference between the BIC for the alternative hypothesis, *BIC*_*H1*_, and the BIC for the null hypothesis, BIC_H0_, defined as

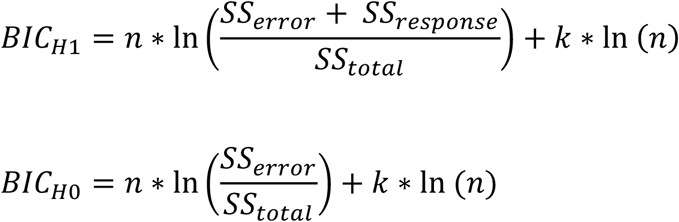

Where *k* is the number of model parameters and *n* is the total sample size. We set *k* equal to two for the models analyzed in this study, comprising the two LMM parameters: intercept and slope

## Acronyms

HFOs: High-frequency oscillations
LFP: local field potential
MUA: multiunit activity
MEA: microelectrode array
BHF: broadband high frequency
MTL: mesial temporal lobe

## SUPPLEMENTARY FIGURE LEGENDS

**Supplementary Figure 1.**
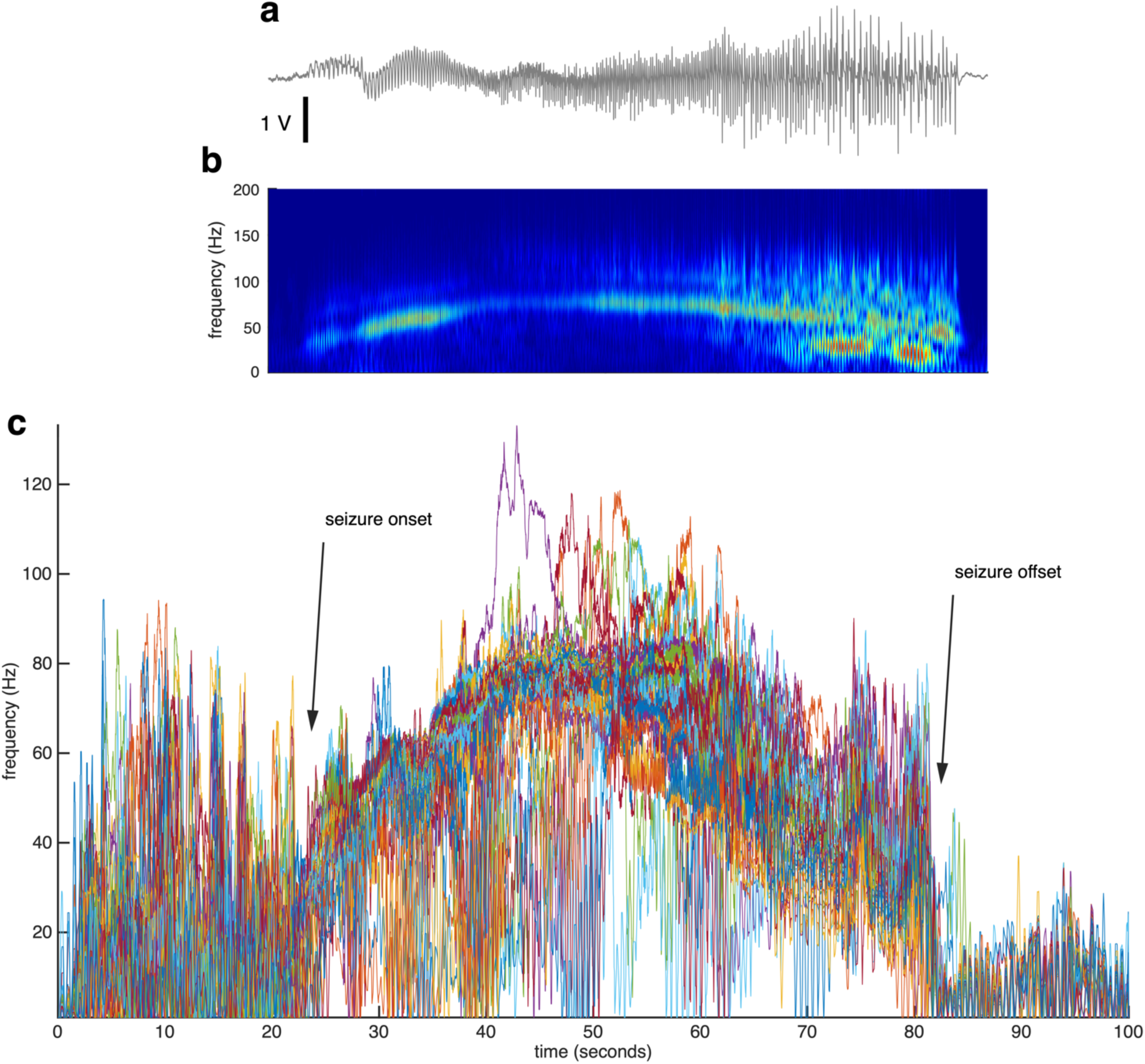
Dominant Frequency Detection. A, example broadband LFP from an ECoG electrode. B, spectrogram of the seizure recorded on the electrode shown in A. C, dominant frequencies for all recorded ECoG electrodes through the duration of the seizure. Each colored line represents the dominant frequency from a distinct ECoG electrode. Seizure onset and offset are indicated with arrows.

**Supplementary Figure 2.**
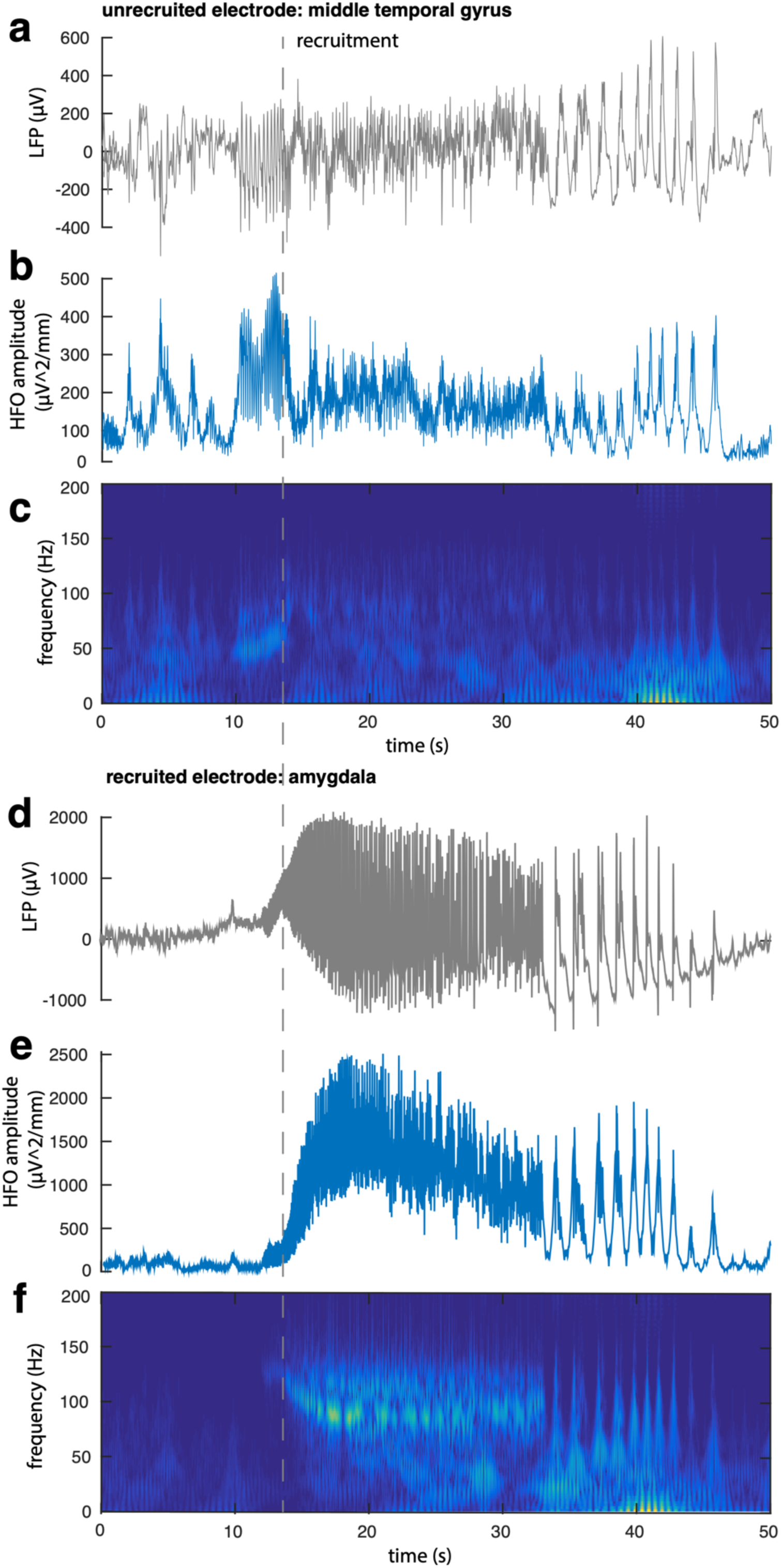
Spatiotemporal profile of recruitment of a seizure in which the core is in the MTL. A, example broadband LFP from an ECoG electrode that is not recruited into the ictal core, i.e. that remains in the penumbra. B, HFO amplitude from the electrode shown in A exhibiting large discharges just before recruitment (the time of recruitment is indicated with gray dotted line). C, A spectrogram of the seizure recorded on the electrode shown in A. D, example broadband LFP from an ECoG electrode that is rapidly recruited into the seizure core. E, HFO amplitude from the electrode shown in D. F, spectrogram of the seizure recorded on the electrode shown in D.

**Supplementary Figure 3.**
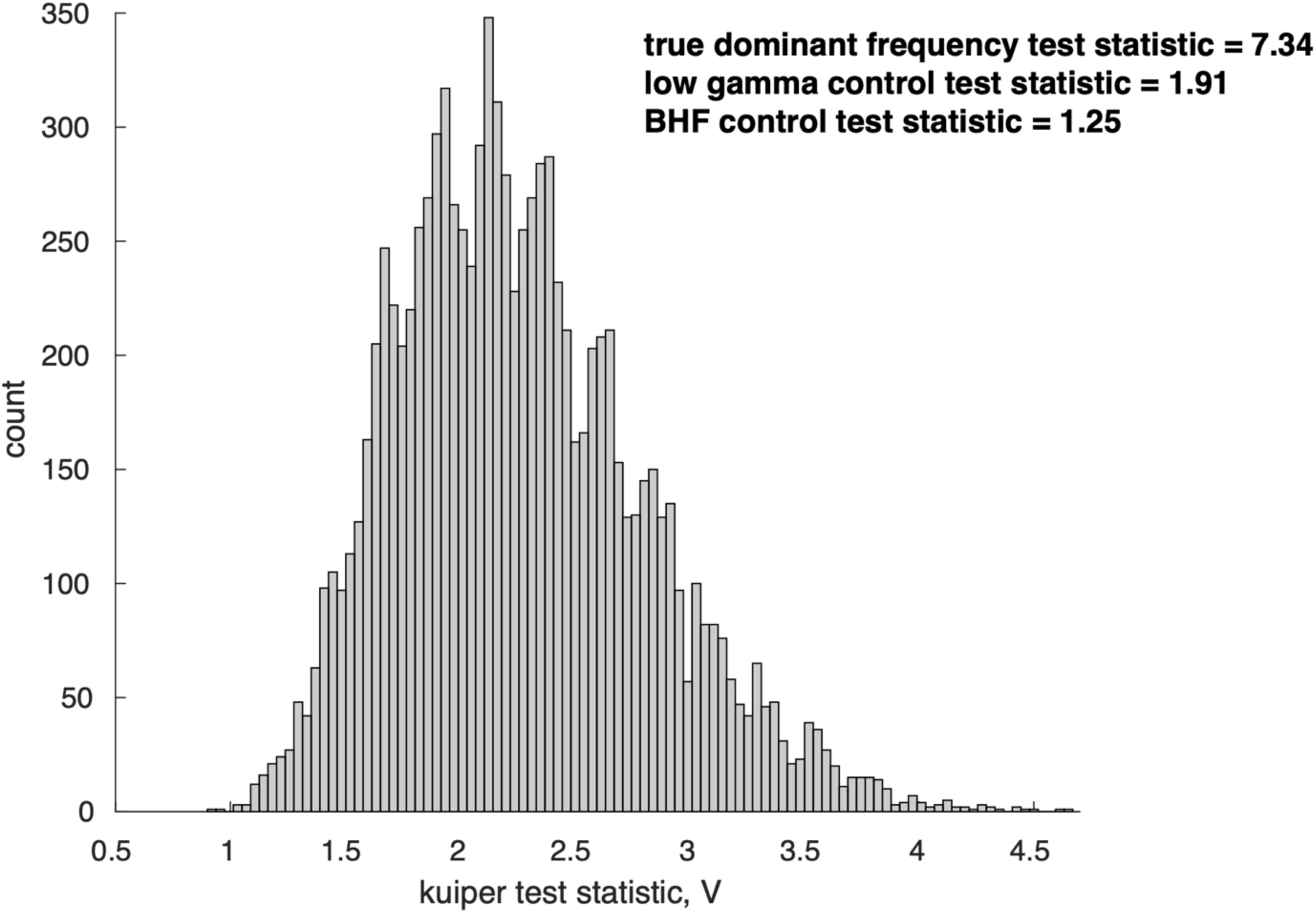
Permutation distribution for Kuiper tests for MUA-HFO coupling. A histogram of test statistics from the shuffled Kuiper tests. The values of test statistics for three analyses performed with the real data from three frequency bands are shown above the histogram.

## REFERENCES

Alfonsa H, Merricks EM, Codadu NK, Cunningham MO, Deisseroth K, Racca C, Trevelyan AJ. 2015. The Contribution of Raised Intraneuronal Chloride to Epileptic Network Activity. J Neurosci 35:7715–7726. doi:10.1523/JNEUROSCI.4105-14.2015

Alvarado-Rojas C, Huberfeld G, Baulac M, Clemenceau S, Charpier S, Miles R, de la Prida LM, Le Van Quyen M. 2015. Different mechanisms of ripple-like oscillations in the human epileptic subiculum. Ann Neurol 77:281–290. doi:10.1002/ana.24324

Bazelot M, Teleńczuk MT, Miles R. 2016. Single CA3 pyramidal cells trigger sharp waves in vitro by exciting interneurones. J Physiol 594:2565–2577. doi:10.1113/JP271644

Bragin A, Mody I, Wilson CL, Jr JE. 2002. Local Generation of Fast Ripples in Epileptic Brain 22:10. doi:https://doi.org/10.1523/JNEUROSCI.22-05-02012.2002

Buzsáki G, Anastassiou CA, Koch C. 2012. The origin of extracellular fields and currents — EEG, ECoG, LFP and spikes. Nat Rev Neurosci 13:407–420. doi:10.1038/nrn3241

Buzsaki G, Horvath Z, Urioste R, Hetke J, Wise K. 1992. High-frequency network oscillation in the hippocampus. Science 256:1025–1027. doi:10.1126/science.1589772

Eissa TL, Tryba AK, Marcuccilli CJ, Ben-Mabrouk F, Smith EH, Lew SM, Goodman RR, McKhann GM, Frim DM, Pesce LL, Kohrman MH, Emerson RG, Schevon CA, van Drongelen W. 2016. Multiscale Aspects of Generation of High-Gamma Activity during Seizures in Human Neocortex. eNeuro 3. doi:10.1523/ENEURO.0141-15.2016

Fedele T, Burnos S, Boran E, Krayenbühl N, Hilfiker P, Grunwald T, Sarnthein J. 2017. Resection of high frequency oscillations predicts seizure outcome in the individual patient. Sci Rep 7:13836. doi:10.1038/s41598-017-13064-1

Foffani G, Uzcategui YG, Gal B, Menendez de la Prida L. 2007. Reduced Spike-Timing Reliability Correlates with the Emergence of Fast Ripples in the Rat Epileptic Hippocampus. Neuron 55:930–941. doi:10.1016/j.neuron.2007.07.040

Freund TF, Katona I. 2007. Perisomatic Inhibition. Neuron 56:33–42. doi:10.1016/j.neuron.2007.09.012

Fried I, Wilson CL, Maidment NT, Engel J, Behnke E, Fields TA, Macdonald KA, Morrow JW, Ackerson L. 1999. Cerebral microdialysis combined with single-neuron and electroencephalographic recording in neurosurgical patients: Technical note. J Neurosurg 91:697–705. doi:10.3171/jns.1999.91.4.0697

Haller M, Donoghue T, Peterson E, Varma P, Sebastian P, Gao R, Noto T, Knight RT, Shestyuk A, Voytek B. 2018. Parameterizing neural power spectra. bioRxiv. doi:10.1101/299859

Höller Y, Kutil R, Klaffenböck L, Thomschewski A, Höller PM, Bathke AC, Jacobs J, Taylor AC, Nardone R, Trinka E. 2015. High-frequency oscillations in epilepsy and surgical outcome. A meta-analysis. Front Hum Neurosci 9. doi:10.3389/fnhum.2015.00574

House PA, MacDonald JD, Tresco PA, Normann RA. 2006. Acute microelectrode array implantation into human neocortex: preliminary technique and histological considerations. Neurosurg Focus 20:1–4.

Ibarz JM, Foffani G, Cid E, Inostroza M, Menendez de la Prida L. 2010. Emergent Dynamics of Fast Ripples in the Epileptic Hippocampus. J Neurosci 30:16249–16261. doi:10.1523/JNEUROSCI.3357-10.2010

Isaacson JS, Scanziani M. 2011. How Inhibition Shapes Cortical Activity. Neuron 72:231–243. doi:10.1016/j.neuron.2011.09.027

Jacobs J, Staba R, Asano E, Otsubo H, Wu JY, Zijlmans M, Mohamed I, Kahane P, Dubeau F, Navarro V, Gotman J. 2012. High-frequency oscillations (HFOs) in clinical epilepsy. Prog Neurobiol 98:302–315. doi:10.1016/j.pneurobio.2012.03.001

Jacobs J, Wu JY, Perucca P, Zelmann R, Mader M, Dubeau F, Mathern GW, Schulze-Bonhage A, Gotman J. 2018. Removing high-frequency oscillations: A prospective multicenter study on seizure outcome. Neurology 91:e1040–e1052. doi:10.1212/WNL.0000000000006158

Jarosz AF, Wiley J. 2014. What Are the Odds? A Practical Guide to Computing and Reporting Bayes Factors. J Probl Solving 7. doi:10.7771/1932-6246.1167

Jefferys JGR, Menendez de la Prida L, Wendling F, Bragin A, Avoli M, Timofeev I, Lopes da Silva FH. 2012. Mechanisms of physiological and epileptic HFO generation. Prog Neurobiol 98:250–264. doi:10.1016/j.pneurobio.2012.02.005

Jirsch JD. 2006. High-frequency oscillations during human focal seizures. Brain 129:1593–1608. doi:10.1093/brain/awl085

Koziol J. A. 1996. A weighted Kuiper statistic for goodness of fit. Stat Neerlandica 50:394–403. doi:10.1111/j.1467-9574.1996.tb01505.x

Manning JR, Jacobs J, Fried I, Kahana MJ. 2009. Broadband Shifts in Local Field Potential Power Spectra Are Correlated with Single-Neuron Spiking in Humans. J Neurosci 29:13613. doi:10.1523/JNEUROSCI.2041-09.2009

Menendez de la Prida L, Huberfeld G, Cohen I, Miles R. 2006. Threshold Behavior in the Initiation of Hippocampal Population Bursts. Neuron 49:131–142. doi:10.1016/j.neuron.2005.10.034

Menendez de la Prida L, Trevelyan AJ. 2011. Cellular mechanisms of high frequency oscillations in epilepsy: On the diverse sources of pathological activities. Epilepsy Res 97:308–317. doi:10.1016/j.eplepsyres.2011.02.009

Merricks EM, Smith EH, McKhann GM, Goodman RR, Bateman LM, Emerson RG, Schevon CA, Trevelyan AJ. 2015. Single unit action potentials in humans and the effect of seizure activity. Brain 138:2891–2906. doi:10.1093/brain/awv208

Miller KJ. 2010. Broadband Spectral Change: Evidence for a Macroscale Correlate of Population Firing Rate? J Neurosci 30:6477. doi:10.1523/JNEUROSCI.6401-09.2010

Miri ML, Vinck M, Pant R, Cardin JA. 2018. Altered hippocampal interneuron activity precedes ictal onset. eLife 7:e40750. doi:10.7554/eLife.40750

Misra A, Burke J, Ramayya A, Jacobs J, Sperling M, Moxon K, Kahana M, Evans J, Sharan A. 2014. Methods for implantation of micro-wire bundles and optimization of single/multiunit recordings from human mesial temporal lobe. J Neural Eng 11:026013. doi:10.1088/1741-2560/11/2/026013

Ray S, Maunsell JHR. 2011. Different Origins of Gamma Rhythm and High-Gamma Activity in Macaque Visual Cortex. PLoS Biol 9:e1000610. doi:10.1371/journal.pbio.1000610

Roux L, Buzsáki G. 2015. Tasks for inhibitory interneurons in intact brain circuits. Neuropharmacology 0:10–23. doi:10.1016/j.neuropharm.2014.09.011

Schevon CA, Weiss SA, McKhann G, Goodman RR, Yuste R, Emerson RG, Trevelyan AJ. 2012a. Evidence of an inhibitory restraint of seizure activity in humans. Nat Commun 3:1060. doi:10.1038/ncomms2056

Schevon CA, Weiss SA, McKhann Jr G, Goodman RR, Yuste R, Emerson RG, Trevelyan AJ. 2012b. Evidence of an inhibitory restraint of seizure activity in humans. Nat Commun 3:1060.

Schlingloff D, Káli S, Freund TF, Hájos N, Gulyás AI. 2014. Mechanisms of Sharp Wave Initiation and Ripple Generation. J Neurosci 34:11385–11398. doi:10.1523/JNEUROSCI.0867-14.2014

Smith EH, Liou J, Davis TS, Merricks EM, Kellis SS, Weiss SA, Greger B, House PA, McKhann II GM, Goodman RR, Emerson RG, Bateman LM, Trevelyan AJ, Schevon CA. 2016. The ictal wavefront is the spatiotemporal source of discharges during spontaneous human seizures. Nat Commun 7:11098.

Trevelyan AJ. 2009. The Direct Relationship between Inhibitory Currents and Local Field Potentials. J Neurosci 29:15299–15307. doi:10.1523/JNEUROSCI.2019-09.2009

Trevelyan AJ, Sussillo D, Yuste R. 2007. Feedforward Inhibition Contributes to the Control of Epileptiform Propagation Speed. J Neurosci 27:3383–3387. doi:10.1523/JNEUROSCI.0145-07.2007

Valero M, Cid E, Averkin RG, Aguilar J, Sanchez-Aguilera A, Viney TJ, Gomez-Dominguez D, Bellistri E, de la Prida LM. 2015. Determinants of different deep and superficial CA1 pyramidal cell dynamics during sharp-wave ripples. Nat Neurosci 18:1281–1290. doi:10.1038/nn.4074

Weiss SA, Banks GP, McKhann GM, Goodman RR, Emerson RG, Trevelyan AJ, Schevon CA. 2013. Ictal high frequency oscillations distinguish two types of seizure territories in humans. Brain 136:3796–3808. doi:10.1093/brain/awt276

Weiss SA, Lemesiou A, Connors R, Banks GP, McKhann GM, Goodman RR, Zhao B, Filippi CG, Nowell M, Rodionov R. 2015. Seizure localization using ictal phase-locked high gamma A retrospective surgical outcome study. Neurology 84:2320–2328.

